# Butterfly abundance declines over 20 years of systematic monitoring in Ohio, USA

**DOI:** 10.1101/613786

**Authors:** Tyson Wepprich, Jeffrey R. Adrion, Leslie Ries, Jerome Wiedmann, Nick M. Haddad

## Abstract

Severe insect declines make headlines, but they are rarely based on systematic monitoring outside of Europe. We estimate the rate of change in total butterfly abundance and the population trends for 81 species using 21 years of systematic monitoring in Ohio, USA. Total abundance is declining at 2% per year, resulting in a cumulative 33% reduction in butterfly abundance. Three times as many species have negative population trends compared to positive trends. The rate of total decline and the proportion of species in decline mirror those documented in three comparable long-term European monitoring programs. Multiple environmental changes such as climate change, habitat degradation, and agricultural practices may contribute to these declines in Ohio and shift the makeup of the butterfly community by benefiting some species over others. Our analysis of life-history traits associated with population trends shows an impact of climate change, as species with northern distributions and fewer annual generations declined more rapidly. However, even common and invasive species associated with human-dominated landscapes are declining, suggesting widespread environmental causes for these trends. Declines in common species, although they may not be close to extinction, will have an outsized impact on the ecosystem services provided by insects. These results from the most extensive, systematic insect monitoring program in North America demonstrate an ongoing defaunation in butterflies that on an annual scale might be imperceptible, but cumulatively has reduced butterfly numbers by a third over 20 years.

## Introduction

Defaunation, or the drastic loss of animal species and declines in abundance, threatens to destabilize ecosystem functioning globally (1). In comparison to studies of vertebrate populations, monitoring of changes in insect diversity is more difficult and far less prevalent (2,3). Despite this, a global analysis of long-term population trends across 452 species estimated that insect abundance had declined 45% over 40 years (1). Recently, more extreme declines in insect biomass have been observed upon resampling after 2-4 decades (4,5). Losses of total biomass or total abundance across all species may be more consequential than local declines in species diversity, as common insect species contribute the most to ecosystem services, such as pollination (6). However, our knowledge of insect declines is skewed towards European monitoring programs, including in global analyses (1). In this study, we analyze long-term, region-wide trends in abundance across a diversity of species for an entire insect group in North America to examine the scope of insect defaunation. The best source of data to assess insect defaunation comes from large-scale, systematic monitoring programs of multiple species (3). Through these efforts, trained volunteers or citizen scientists have contributed much of the evidence for biotic responses to anthropogenic climate warming such as changes in insect phenology and range distributions (7,8). Unlike citizen science reporting of opportunistic observations or species checklists, many insect monitoring programs use a systematic protocol developed specifically to track butterfly abundances through time, both within and between seasons, and over large spatial scales (9). Pollard-based monitoring programs, modeled after the first nationwide Butterfly Monitoring Scheme launched in the United Kingdom in 1977 (UKBMS), use weekly standardized counts on fixed transects (10). Their widespread adoption enables regional comparisons of insect responses to environmental change or defaunation (11,12). We compare our analysis with exemplary long-term monitoring schemes from Europe to test if the rate of insect declines generalizes across continents.

The best source of abundance data for assessment of chronic insect decline, and the most prominent source of data in (1), is within the butterflies. Due to the relative ease and popularity of monitoring butterflies, environmental assessments use them as an indicator taxa for the general trajectory of biodiversity, assuming that they experience comparable pressures from land-use change, climate change, and habitat degradation as other insect taxa (13–15). Intensive long-term monitoring of individual butterfly species has provided rigorous, quantitative estimates of declines. Most prominently, the Eastern North American Monarch has declined by over 85% (16) and the Western North American Monarch by over 95% (17) over the past two decades. Severe declines have also been observed in some of the rarest butterflies (18,19). These data from individual species of conservation concern may not represent a broader trend across butterflies, which is what we aim to document in this study. Volunteers, organized and trained by The Ohio Lepidopterists, have assembled the most extensive dataset of systematic butterfly counts that stands alone in North America in terms of the spatial extent and sampling frequency of Pollard walks (9). Three other monitoring programs in the United States have documented long-term, multi-species population trends. In Massachusetts, based on species lists from field trips, climate-driven community shifts explain how the relative likelihood of species observations change over 18 years (20). Shapiro and colleagues have made biweekly presence/absence observations and Pollard-based counts on 11 fixed transects along an elevational gradient in California over more than 45 years to document species richness changes in response to climate and land-use, increasing abundance at a high elevation site, and impacts of agricultural practices on abundance at low elevation sites (21,22). Several teams have monitored declines in specialist butterflies restricted to native prairie patches in the Midwestern states with transect or timed survey methods over 26 years (23,24). The growing number of Pollard-based monitoring programs in the United States (9) has the potential to track how widespread and consistent butterfly trends are across regions. Here, we used 21 years of weekly butterfly surveys across 104 sites to assess abundance trends for butterflies in Ohio. We estimate population trends for 81 species and test for their association with life-history traits and phylogenetic relatedness. We review findings from European butterfly monitoring schemes for quantitative comparison with the rate of abundance changes in Ohio. This analysis provides evidence of widespread insect defaunation and species’ declines from the most extensive, systematic monitoring program in North America.

## Materials and methods

### Study sites

We studied butterfly population trends across the state of Ohio in the Midwestern USA. Over its 116,100 km^2^ land area, Ohio has a mosaic of habitat types due to its partially glaciated history and its place at the confluence of Midwestern prairies, the Appalachian Mountains, and the boreal forest (25). Only remnants of wetland and prairie habitat remain in the state due to human modification of the landscape. Some rare butterflies have declined due to forest succession following suppression of disturbances (26). Agriculture and pastures (50%), forest (30%), and urban development (10%) are the predominant land-use/land cover classes (27).

Monitoring sites have a Northeast to Southwest gradient in their mean annual temperatures (mean 18.8°C, range from 14.0°C to 23.6°C) from interpolated daily temperatures from Daymet over 1996-2016 (Thornton et al. 1997). Mean annual temperatures at these sites grew at a linear trend of 0.3°C per decade and growing season length has increased by 60 degree-days (base 5°C) per decade from 1980-2016. Monitoring sites span the state but are concentrated near cities (Fig 1). On average, within a radius of 2 kilometers, monitoring sites have 24% cropland and pasture, 34% forest, and 30% urban land-use based on the National Land Cover Dataset (29). Although not considered in this study, impervious surfaces from urban development influence temperature-dependent butterfly phenology in Ohio through the urban heat island effect, which may not be fully captured in these gridded temperature interpolations (30).

**Fig 1:**
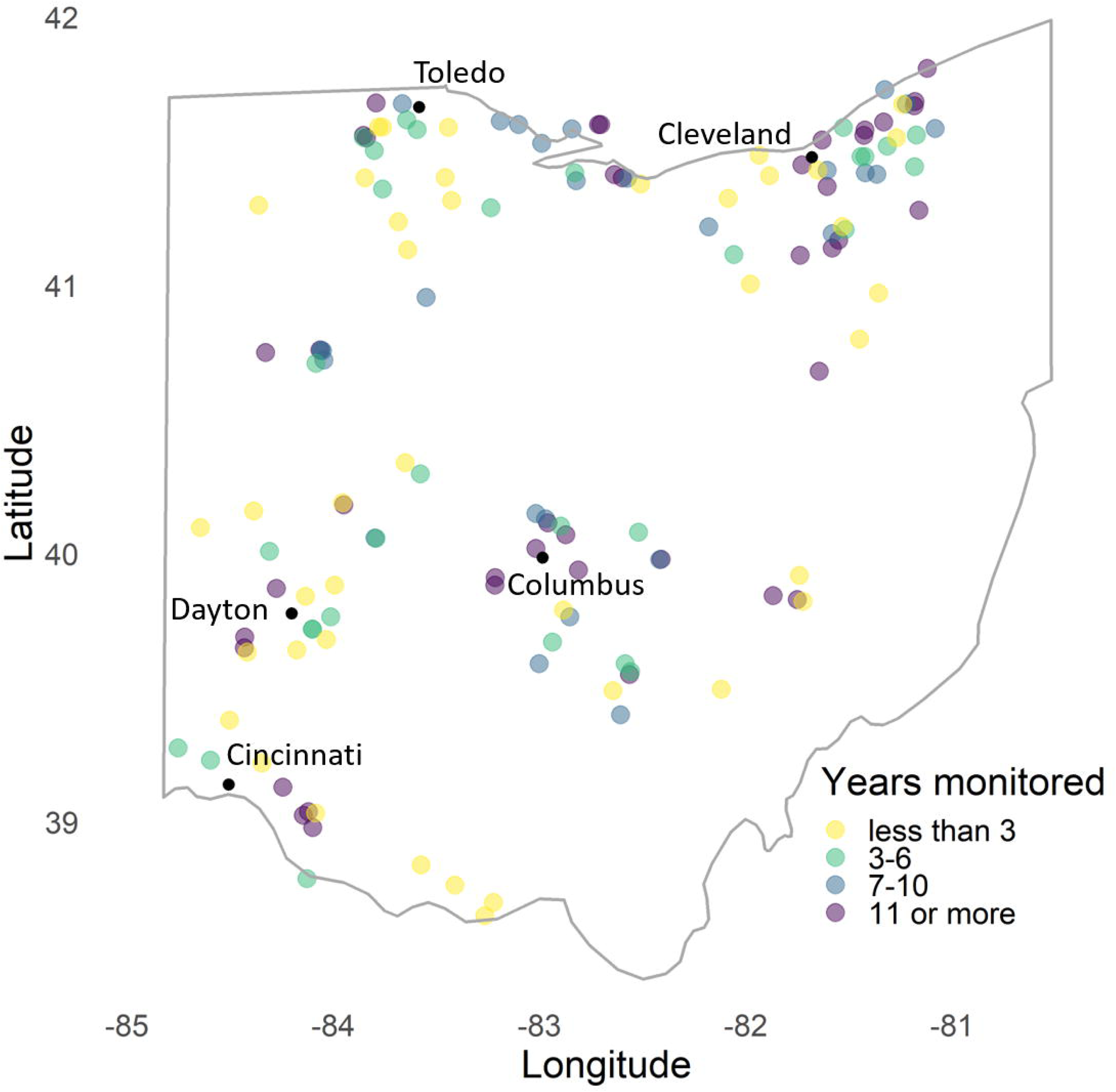
Transect locations monitored by volunteers with the Ohio Lepidopterists. Of the 147 sites, this analysis used the 104 sites monitored for three or more years.

### Monitoring surveys

Trained volunteers contributed 24,405 butterfly surveys from 1996 to 2016 as part of the Ohio Lepidopterists Long-term Monitoring of Butterflies program. Volunteers surveyed on fixed paths at approximately weekly intervals during the entire growing season from April through October (median 23 of 30 weeks surveyed per year per site) and count every species within an approximate 5-meter buffer around the observer (10). Surveys are constrained to times of good weather to increase the detectability of butterflies and last a mean 85 minutes in duration. The annual number of monitored sites ranged from 13 in 1996 to a maximum of 80 in 2012. We limited our analysis of abundance trends to the 104 sites with three or more years of monitoring data and 10 or more surveys per year at each site (Fig 1). We included observations of all sites with at least 5 surveys per year in phenology models that we used to interpolate missing counts before estimating abundance (31).

All 102 species with population indices estimated by phenology models contributed to the total abundance analysis. We limited species-specific analysis to 81 species with sufficient population indices for estimating trends (present at five or more sites and for 10 or more years). Species naming conventions in the monitoring program follow those used in Opler and Krizek (1984) and Iftner et al. (1992) except for combining all observations of *Celastrina ladon* (Spring Azure) and *Celastrina neglecta* (Summer Azure) as an unresolved species complex.

### Population indices

We estimated population indices for each site x year x species by adapting methods established for the UKBMS that account for missing surveys and butterfly phenology over the season (31,33). We used generalized additive models for each species to estimate variation in counts in order to interpolate missing surveys with model predictions (31,34). To account for seasonal, spatial, and interannual variation in species phenology, we extended the regional generalized additive model approach (12, Supplement 1) by including spatially-explicit site locations and converting calendar dates of observations to degree-days (35), which can improve butterfly phenology predictions (36). We calculated the population index by integrating over the weekly counts and missing survey interpolations using the trapezoid method (31).

### Controlling for confounding factors

We accounted for differences in sampling across sites and years so that our modeled trends would capture changes in abundance rather than changes in detection probability (37). True abundance is confounded with detection probability when using counts from Pollard walks (38). Butterfly monitoring protocols that account for detection probability like distance sampling are commonly used for single-species studies (39), but untenable for scaling up to a regional program. Most analyses of Pollard walks assume no systematic change in detectability (but see (40)) because counts correlate closely with true abundance estimates from distance sampling (41,42). We used two covariates to account for variation in sampling and its influence on population indices for each site x year (20,37,43). We tracked the mean number of species reported in each survey, or list-length, which is a synthetic measure of factors influencing detectability such as weather conditions, site quality, and observer effort (20,44,45). We treated the total duration of surveys in minutes as an offset in the models of population trends. Because we interpolated missing surveys for the population indices, we projected what the total duration would be if all 30 weeks had been surveyed at the mean duration reported for that site x year.

Sampling across the state is nonrandom because participants choose transect locations, a common practice in volunteer-based monitoring programs. Since sites generally cluster near human population centers with a greater proportion of developed land-use and a lesser proportion of agriculture, we assumed that population trends at the 104 sites across the state sufficiently capture the broader statewide trends (37). Comparisons between the UKBMS volunteer-placed transects and a broader survey with stratified, random sampling show congruence between species trends estimated from each monitoring strategy (46).

### Population Trends

We used generalized linear mixed models to estimate temporal trends in relative abundance for 81 species from their population indices (47). We modeled population indices at each site and year as an over-dispersed Poisson random variable with covariates on the log-link scale.

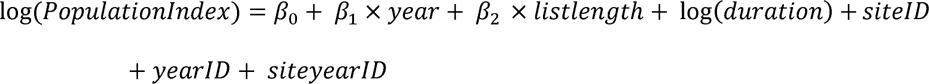

We included the numeric year and mean list length for each population index as covariates, which were centered to aid in model fitting and interpretation (48). We used the coefficient for year (β_2_) as the annual trend in population indices as our main result. We controlled for changes in sampling by using the total duration of surveys as a model offset, converting the dependent variable to a rate of butterflies counted per minute. Random effects of individual sites and years account for spatial and temporal variation in population counts deviating from the statewide trend. We accounted for over-dispersion in the Poisson-distributed counts with the random effect *siteyearID* for each unique observation (49). We modeled trends in total abundance using the same modeling approach, but summed across 102 species’ population indices for each site x year observation. We interpreted trends as an annual rate by taking the geometric mean rate of change between the predicted abundance between two points in time after setting the list-length covariate to its mean and excluding the random effects (47). For comparisons with other monitoring programs, we used a *p*-value threshold of 0.05 to classify trends as positive, negative, or stable.

Our approach is similar to that used by the UKBMS and other European monitoring programs which use generalized linear models in TRIM software (50). One key difference is that our site and annual fluctuations from the temporal trend were derived from random effects rather than fixed effects, which reduces spurious detection of trends (43). Another key difference is that TRIM does not allow for continuous covariates, which we used to account for sampling variation instead of assuming no confounding pattern in sampling effort. To validate that our modeling choices did not unreasonably influence the results, we used three alternative approaches: (1) a Poisson-based generalized linear model (equation 1 without the random effect *siteyearID*); (2) a nonlinear generalized additive mixed model with a smoothing spline replacing the linear temporal trend (43); and (3) a TRIM model with over-dispersion and serial temporal correlation but no sampling covariates or offsets (50). We compared similarity in the total abundance trends, the correlation of species’ trends between model alternatives, and the classification of species’ trends as positive, stable, or negative.

### Comparison with other studies

We compare our findings to three European long-term, regional butterfly monitoring programs with systematic Pollard walks that publish regular updates on total abundance and species’ trends (40,51,52). Although all programs analyzed counts with Poisson regression, we had to standardize them differently depending on the data available and their modeling approaches. The UKBMS reports total abundance indicators as the geometric mean of species trends from two groups: specialist and countryside species (51). We used the reported smoothed annual index values for these indicators because the first year of monitoring is an outlier that exaggerates declines (UK Biodiversity Indicators 2018, http://jncc.defra.gov.uk/page-4236). We used the Dutch Butterfly Monitoring Scheme’s reported cumulative annual trend in total butterflies counted across all transects after correction for missing surveys (52). For the Catalan Butterfly Monitoring Scheme, we extracted annual population indices from the 2015-2016 annual report (53) with WebPlotDigitizer 4.1 (54) and performed a Poisson regression over time with annual random effects to obtain a comparable abundance trend. We converted total abundance trends into annual percent rates for comparison. We tallied the increases and decreases in species’ trends for each region reported by the monitoring program, without accounting for differences in their statistical approaches.

### Species’ traits

To explore potential mechanisms that might explain species-level variation in abundance trends, we modeled the estimates of species’ temporal trends (/3_1_) as a response to life history traits (20,30). Of the 81 species, we classified 14 as migratory species and 67 as year-round residents of Ohio. We analyzed traits models both across all species and after excluding migratory species, which would have population trends driven by factors outside of Ohio. We collected traits that relate to insect responses to climate change and habitat change, as these are two primary drivers of butterfly community changes (7,20,21).

We tested if butterflies with traits making them more adaptive to a warming climate have more positive population trends. We compared species with different range distributions, assuming that species distributed in warmer, Southern regions would be more likely to increase in Ohio as the climate warms. We assigned species’ ranges as Southern, core, or Northern by range maps and county records (25,32). Voltinism, or the number of generations per year, increases in warmer years and warmer regions within many species in Ohio (55), compared with obligate univoltine species that do not adjust their lifecycle based on changing growing season length. We assigned voltinism observed in Ohio as univoltine, bivoltine, or multivoltine (3+ generations per year) based on visualization of phenology models and (25). The life stage in which species overwinter, obtained from (25), contributes to its ability to respond to warming with shifts in phenology (20,56).

We would expect more generalist species, in host plant requirements and habitat preferences, to have more positive population trends in a landscape heavily modified by human use (21,51). For host plant requirements, we gathered two traits from the literature that describe host plant category (forb, graminoid, or woody) and whether the butterfly’s host plant requirements span multiple plant families or are limited to one plant family or genus (25). Mean wing size from (32) was used as a surrogate of dispersal ability between habitats, which is expected to increase ability to access resources in a fragmented landscape. Three of the authors assigned species as wetland-dependent or human-disturbance tolerant species, which we aggregated into two binary variables to test if these specialist or generalist habitat preferences correlate with abundance trends.

We used univariate linear models for each life history trait both for all 81 species and with the 14 migratory species excluded. To account for the phylogenetic relatedness and the non-independence across species, we also used phylogenetic generalized least squares models that estimated branch length transformations with Pagel’s lambda by maximum likelihood (57). The phylogenetic models excluded three species without gene sequences available.

### Phylogenetic tree

We obtained coding sequences for the most widely used DNA barcoding locus, the mitochondrial cytochrome c oxidase subunit I gene COI-5P, from GenBank (58). For species not found in GenBank, we obtained coding sequences from The Barcode of Life Data System (59). When possible, we obtained sequences from multiple sampling locations in North America.

Owing to the relatively small size of our multiple-species alignment—i.e. a single mtDNA locus, 651 base pairs in length—we decided to take both a constrained and unconstrained maximum likelihood approach to estimate the genealogical relationships of our samples. Some of the species from our analysis, though not all, were recently used in a more comprehensive phylogenetic analysis of butterflies (60), thus prompting us to constrain the phylogenetic backbone of our tree using family-level relationships. We report details of our workflow in Supplement 1.

### Statistical analysis

We used R 3.5.2 for analysis (61) and share the data and our code on Dryad. We fit generalized additive models with the *mgcv* package (34), generalized linear mixed models with the *lme4* package (Bates et al. 2015), generalized additive mixed models with the *poptrend* package (43), and phylogenetic generalized least squares models with the *ape* and *caper* packages (63,64). Confidence intervals for the temporal trends were estimated with bootstrapped model fits with the *merTools* and *poptrend* packages (43,65). For models of population trends, we estimated the goodness of fit with *R^2^* developed for generalized linear mixed models that give marginal and conditional *R^2^* values for the fixed effects and the fixed + random effects, respectively (66,67). For trait models, we reported the adjusted *R^2^* values from the univariate models.

## Results

The statewide relative abundance summed across all species declined at an annual rate of 2.0% (*β*_1_= -0.020, std. err. 0.005, *p* < 0.001), accumulating a 33% decline over 1996-2016 (Table 1, Fig 2). Among population trends, more than three times as many species are declining than increasing in abundance at our threshold of *p* < 0.05 (32 versus 9, respectively) (Table 2, Fig 3 for migratory species and Fig 4 for resident species). Positive and negative species trends are distributed across the phylogenetic tree (Fig A in S1 Appendix).

**Fig 2:**
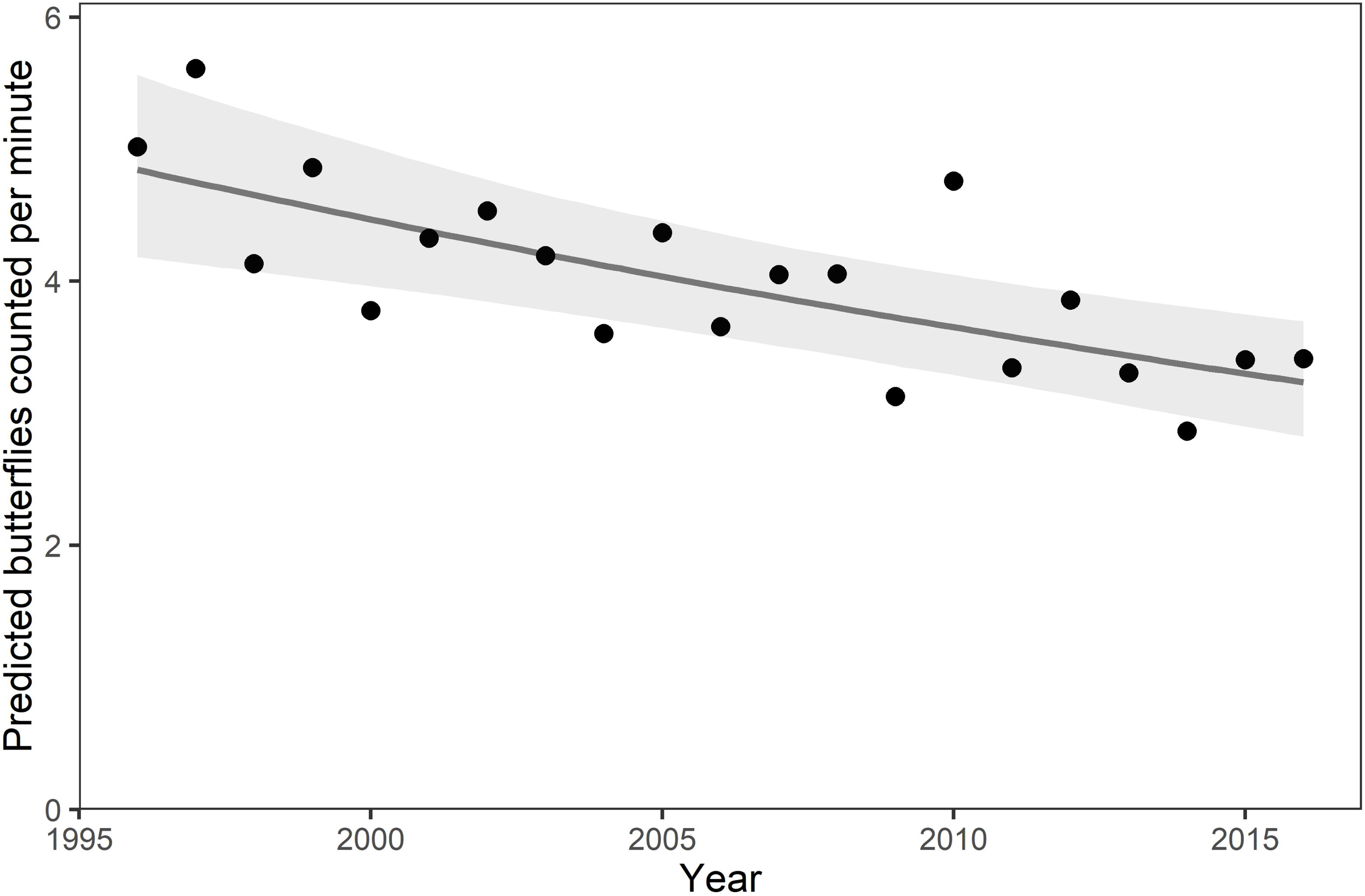
The statewide relative abundance of butterflies (all species aggregated) in Ohio declined by 33% over 1996-2016. Plotted are model predictions for each year based on the fixed effects of year (solid line) and annual random effects (dots) to show annual variation about the trend line. Shading shows the 95% confidence interval based on bootstrapped model fits for the temporal trend.

**Fig 3:**
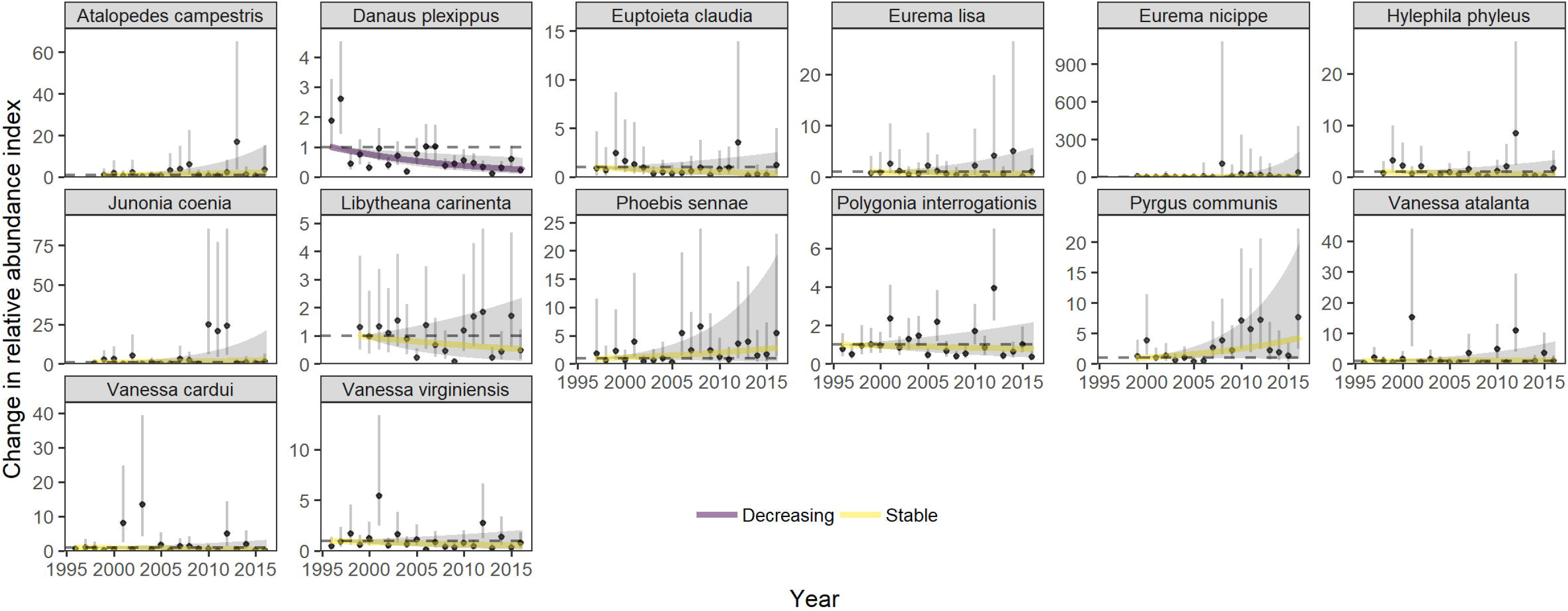
Statewide trends of 14 migratory species with annual variation. Plotted are model predictions for each year based on the fixed effects of year (solid line) and annual random effects (dots) to show annual variation about the trend line. Shading shows 95% confidence intervals based on bootstrapped model fits in the *poptrend* package (43) for the temporal trend and for the annual random effects. The first year’s estimate is set to a value of 1 as a baseline for relative population changes.

**Fig 4:**
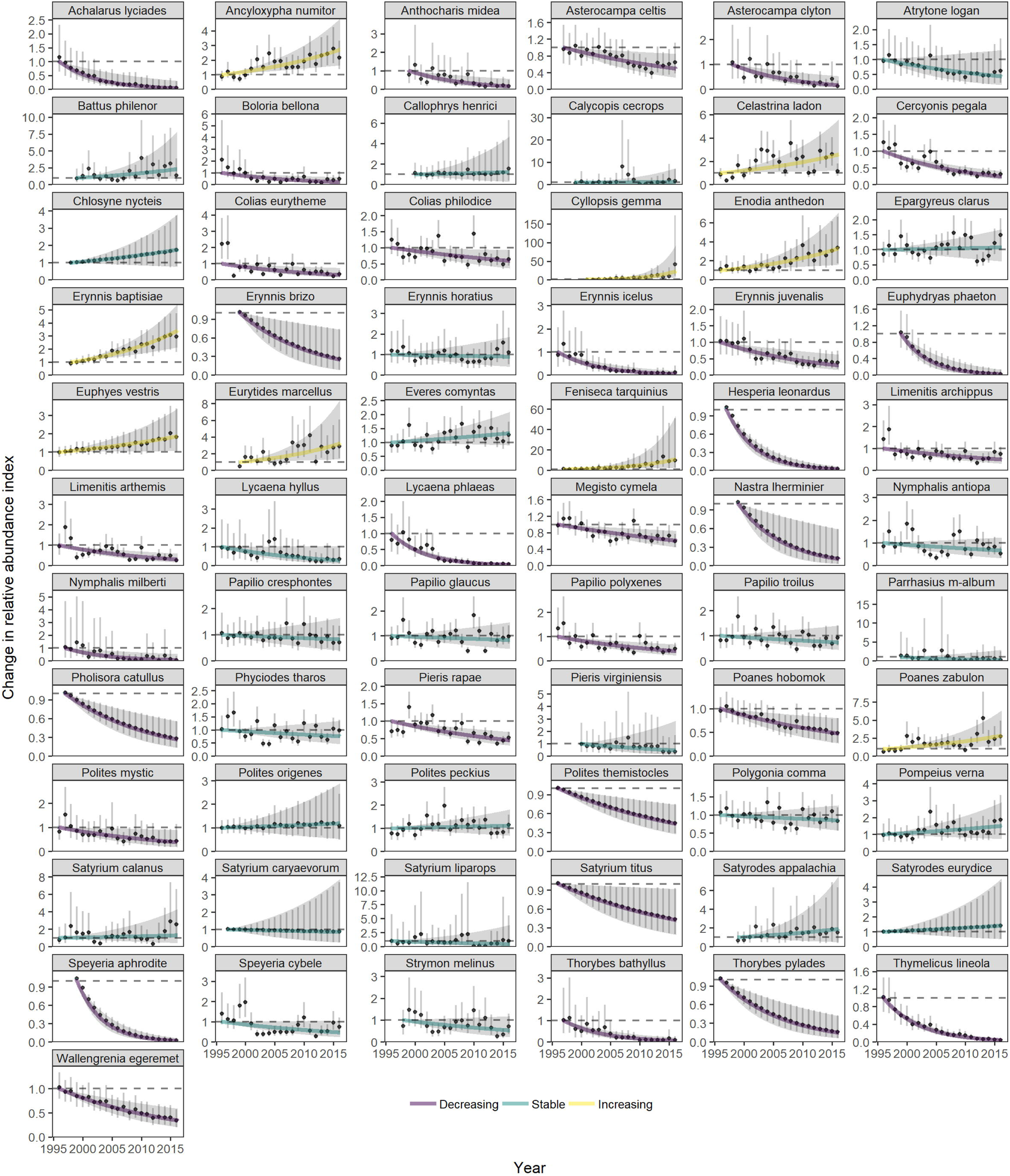
Statewide trends of 67 resident species with annual variation. Plotted are model predictions for each year based on the fixed effects of year (solid line) and annual random effects (dots) to show annual variation about the trend line. Shading shows 95% confidence intervals based on bootstrapped model fits in the *poptrend* package (43) for the temporal trend and for the annual random effects. The first year’s estimate is set to a value of 1 as a baseline for relative population changes.

**Table 1:** Generalized linear mixed model of total abundance across all species. The natural logarithm of the total survey duration across the monitoring season was an offset in the model. The model’s marginal R^2^ was 0.20 for its fixed effects and its conditional R^2^ was 0.61 when including variation in sites, years, and over-dispersion with random effects parameters.

| Fixed effects | *B* | std.error | z statistic | p.value |
| --- | --- | --- | --- | --- |
| Intercept | 1.33 | 0.0506 | 26.4 | <0.001 |
| Year (numeric) | -0.0203 | 0.00496 | -4.11 | <0.001 |
| List-length | 0.104 | 0.00587 | 17.7 | <0.001 |
| Random effects | std. dev. | # groups |  |  |
| Site x year ID | 0.278 | 1005 |  |  |
| Site ID | 0.417 | 104 |  |  |
| Year ID (factor) | 0.121 | 21 |  |  |

**Table 2:** Species’ abundance trends over time. Trends are the coefficient of year in our generalized linear mixed models with the accompanying standard error and *p*-value for the coefficient (equation 1). We show the data available for each species’ model: total number of butterflies recorded for all years, number of sites, number of years, and the number of population indices calculated for each species for use in abundance model (Site x year). Bold font indicates trends that were classified as increasing or decreasing (*p* < 0.05).

| Species |  | Sample size |  |  |  | GLMM temporal trend |  |  |
| --- | --- | --- | --- | --- | --- | --- | --- | --- |
| Common | Latin | Total #<br>counted | Sites | Years | Site/<br>year | Trend<br>coef. | Std.<br>error | P |
| Aphrodite Fritillary | *Speyeria aphrodite* | 477 | 9 | 16 | 131 | **-0.233** | **0.060** | **<0.001** |
| Baltimore | *Euphydryas phaeton* | 818 | 7 | 17 | 83 | **-0.224** | **0.071** | **0.002** |
| American Copper | *Lycaena phlaeas* | 10,255 | 31 | 21 | 359 | **-0.193** | **0.024** | **<0.001** |
| Hoary Edge Skipper | *Achalarus lyciades* | 291 | 7 | 19 | 88 | **-0.178** | **0.061** | **0.003** |
| Milbert's Tortoise Shell | *Nymphalis milberti* | 140 | 8 | 16 | 101 | **-0.174** | **0.065** | **0.008** |
| European Skipper | *Thymelicus lineola* | 46,549 | 57 | 21 | 609 | **-0.173** | **0.021** | **<0.001** |
| Southern Cloudywing | *Thorybes bathyllus* | 667 | 15 | 20 | 194 | **-0.129** | **0.037** | **<0.001** |
| Falcate Orangetip | *Anthocharis midea* | 756 | 8 | 18 | 103 | **-0.123** | **0.040** | **0.002** |
| Dreamy Duskywing | *Erynnis icelus* | 879 | 18 | 21 | 260 | **-0.120** | **0.024** | **<0.001** |
| Swarthy Skipper | *Nastra lherminier* | 448 | 7 | 17 | 78 | **-0.114** | **0.041** | **0.006** |
| Tawny Emperor | *Asterocampa clyton* | 937 | 27 | 19 | 308 | **-0.114** | **0.036** | **0.002** |
| Leonard's Skipper | *Hesperia leonardus* | 1,348 | 11 | 20 | 144 | **-0.110** | **0.025** | **<0.001** |
| White M Hairstreak | *Parrhasius m-album* | 95 | 7 | 15 | 110 | -0.105 | 0.081 | 0.195 |
| Northern Cloudywing | *Thorybes pylades* | 547 | 16 | 20 | 210 | **-0.095** | **0.033** | **0.004** |
| Coral Hairstreak | *Satyrrium titus* | 607 | 15 | 21 | 217 | **-0.094** | **0.025** | **<0.001** |
| Juvenal's Duskywing | *Erynnis juvenalis* | 3,838 | 38 | 21 | 487 | **-0.083** | **0.020** | **<0.001** |
| Common Wood Nymph | *Cercyonis pegala* | 21,603 | 77 | 21 | 788 | **-0.073** | **0.013** | **<0.001** |
| Common Sooty Wing | *Pholisora catullus* | 1,142 | 34 | 20 | 398 | **-0.072** | **0.015** | **<0.001** |
| Sleepy Duskywing | *Erynnis brizo* | 811 | 13 | 18 | 156 | **-0.071** | **0.032** | **0.027** |
| Monarch | *Danaus plexippus* | 46,070 | 104 | 21 | 1,005 | **-0.070** | **0.023** | **0.002** |
| Red-spotted Purple | *Limenitis arthemis* | 6,226 | 87 | 21 | 913 | **-0.064** | **0.019** | **<0.001** |
| Bronze Copper | *Lycaena hyllus* | 656 | 23 | 21 | 254 | -0.063 | 0.039 | 0.103 |
| Northern Broken-Dash | *Wallengrenia egeremet* | 5,959 | 49 | 21 | 528 | **-0.062** | **0.018** | **<0.001** |
| Tawny-edged Skipper | *Polites themistocles* | 2,322 | 48 | 21 | 541 | **-0.058** | **0.016** | **<0.001** |
| West Virginia White | *Pieris virginianensis* | 214 | 5 | 16 | 63 | -0.058 | 0.059 | 0.329 |
| Fiery Skipper | *Hylephila phyleus* | 3,917 | 57 | 19 | 646 | -0.057 | 0.061 | 0.351 |
| Meadow Fritillary | *Boloria bellona* | 5,447 | 55 | 21 | 598 | **-0.056** | **0.027** | **0.040** |
| Orange Sulphur | *Colias eurytheme* | 62,160 | 101 | 21 | 996 | **-0.055** | **0.021** | **0.008** |
| Long Dash | *Polites mystic* | 1,317 | 21 | 21 | 219 | **-0.047** | **0.020** | **0.022** |
| American Lady | *Vanessa virginiensis* | 2,029 | 54 | 21 | 637 | -0.045 | 0.033 | 0.179 |
| Black Swallowtail | *Papilio polyxenes* | 12,410 | 92 | 21 | 941 | **-0.044** | **0.015** | **0.004** |
| Gray Hairstreak | *Strymon melinus* | 2,418 | 49 | 19 | 587 | -0.044 | 0.026 | 0.089 |
| Painted Lady | *Vanessa cardui* | 5,564 | 80 | 21 | 873 | -0.042 | 0.054 | 0.440 |
| Great Spangled Fritillary | *Speyeria cybele* | 33,573 | 90 | 21 | 904 | **-0.041** | **0.020** | **0.047** |
| Hobomok Skipper | *Poanes hobomok* | 6,863 | 51 | 21 | 576 | **-0.040** | **0.014** | **0.005** |
| Viceroy | *Limnitis archippus* | 16,079 | 85 | 21 | 896 | **-0.039** | **0.016** | **0.014** |
| Cabbage White | *Pieris rapae* | 304,105 | 104 | 21 | 1,005 | **-0.038** | **0.010** | **<0.001** |
| Hackberry Emperor | *Asterocampa celtis* | 9,992 | 42 | 20 | 467 | **-0.037** | **0.017** | **0.033** |
| Striped Hairstreak | *Satyrrium liparops* | 155 | 14 | 18 | 211 | -0.028 | 0.067 | 0.682 |
| Variegated Fritillary | *Euptoieta claudia* | 956 | 17 | 19 | 204 | -0.027 | 0.052 | 0.603 |
| Little Wood Satyr | *Megisto cymela* | 76,612 | 87 | 21 | 878 | **-0.026** | **0.009** | **0.005** |
| American Snout Butterfly | *Libytheana carinenta* | 1,007 | 36 | 18 | 418 | -0.025 | 0.050 | 0.612 |
| Hickory Hairstreak | *Satyrrium caryaevorum* | 196 | 12 | 20 | 170 | -0.023 | 0.053 | 0.656 |
| Mourning Cloak | *Nymphalis antiopa* | 3,214 | 85 | 21 | 905 | -0.021 | 0.018 | 0.256 |
| Clouded Sulphur | *Colias philodice* | 49,267 | 102 | 21 | 998 | -0.014 | 0.014 | 0.286 |
| Spicebush Swallowtail | *Papilio troilus* | 25,322 | 82 | 21 | 858 | -0.014 | 0.014 | 0.324 |
| Dun Skipper | *Euphyes vestris* | 1,684 | 49 | 21 | 585 | -0.014 | 0.012 | 0.224 |
| Question Mark | *Polygonia interrogationis* | 6,564 | 88 | 21 | 915 | -0.012 | 0.025 | 0.640 |
| Delaware Skipper | *Atrytone logan* | 1,086 | 30 | 21 | 313 | -0.011 | 0.029 | 0.697 |
| Horace's Duskywing | *Erynnis horatius* | 2,885 | 31 | 21 | 376 | -0.011 | 0.023 | 0.633 |
| Eastern Tiger Swallowtail | *Papilio glaucus* | 29,299 | 101 | 21 | 996 | -0.010 | 0.015 | 0.483 |
| Pearl Crescent | *Phyciodes tharos* | 180,631 | 104 | 21 | 1,005 | -0.010 | 0.014 | 0.461 |
| Little Yellow | *Eurema lisa* | 1,681 | 24 | 18 | 287 | -0.008 | 0.073 | 0.917 |
| Eastern Comma | *Polygonia comma* | 6,222 | 92 | 21 | 944 | -0.007 | 0.011 | 0.561 |
| Giant Swallowtail | *Papilio cresphontes* | 1,109 | 28 | 21 | 322 | 0.002 | 0.019 | 0.912 |
| Banded Hairstreak | *Satyrrium calanus* | 1,107 | 36 | 21 | 468 | 0.004 | 0.031 | 0.896 |
| Silver-spotted Skipper | *Epargyreus clarus* | 54,462 | 102 | 21 | 996 | 0.005 | 0.012 | 0.672 |
| Red Admiral | *Vanessa atalanta* | 28,637 | 97 | 21 | 969 | 0.008 | 0.044 | 0.865 |
| Red-banded Hairstreak | *Calycopis cecrops* | 795 | 7 | 17 | 91 | 0.009 | 0.057 | 0.879 |
| Crossline Skipper | *Polites origenes* | 1,087 | 27 | 21 | 347 | 0.009 | 0.020 | 0.636 |
| Sachem | *Atalopedes campestris* | 1,445 | 19 | 18 | 231 | 0.013 | 0.058 | 0.823 |
| Peck's Skipper | *Polites peckius* | 23,702 | 90 | 21 | 905 | 0.014 | 0.014 | 0.306 |
| Northern Eyed Brown | *Satyroides eurydice* | 1,342 | 13 | 21 | 174 | 0.016 | 0.035 | 0.651 |
| Eastern Tailed Blue | *Everes comyntas* | 56,137 | 99 | 21 | 974 | 0.016 | 0.010 | 0.113 |
| Henry's Elfin | *Callophrys henrici* | 330 | 7 | 17 | 76 | 0.017 | 0.055 | 0.752 |
| Little Glassy Wing | *Pompeius verna* | 8,658 | 56 | 21 | 632 | 0.019 | 0.019 | 0.307 |
| Silvery Checkerspot | *Chlosyne nycteis* | 2,049 | 20 | 19 | 224 | 0.039 | 0.022 | 0.074 |
| Spring/Summer Azure | *Celastrina ladon/neglecta* | 63,947 | 103 | 21 | 1,002 | **0.047** | **0.021** | **0.022** |
| Common Buckeye | *Junonia coenia* | 15,771 | 73 | 19 | 834 | 0.050 | 0.067 | 0.459 |
| Pipevine Swallowtail | *Battus philenor* | 703 | 23 | 18 | 279 | 0.053 | 0.033 | 0.110 |
| Least Skipper | *Ancyloxypha numitor* | 27,506 | 84 | 21 | 844 | **0.053** | **0.015** | **<0.001** |
| Appalachian Eyed Brown | *Satyroides appalachia* | 2,118 | 12 | 18 | 118 | 0.060 | 0.045 | 0.181 |
| Zabulon Skipper | *Poanes zabulon* | 10,960 | 71 | 21 | 747 | **0.061** | **0.022** | **0.004** |
| Northern Pearly-Eye | *Enodia anthedon* | 2,785 | 37 | 21 | 434 | **0.071** | **0.020** | **<0.001** |
| Zebra Swallowtail | *Eurytides marcellus* | 1,349 | 18 | 18 | 224 | **0.075** | **0.030** | **0.011** |
| Cloudless Sulphur | *Phoebis sennae* | 1,840 | 27 | 19 | 355 | 0.088 | 0.057 | 0.121 |
| Common Checkered-Skipper | *Pyrgus communis* | 3,089 | 33 | 18 | 357 | **0.092** | **0.046** | **0.046** |
| Wild Indigo Duskywing | *Erynnis baptisiae* | 15,209 | 51 | 19 | 570 | **0.106** | **0.020** | **<0.001** |
| Harvester | *Feniseca tarquinius* | 341 | 11 | 20 | 143 | **0.122** | **0.061** | **0.046** |
| Sleepy Orange | *Eurema nicippe* | 2,028 | 6 | 17 | 63 | 0.146 | 0.134 | 0.276 |
| Gemmed Satyr | *Cyllopsis gemma* | 1,059 | 6 | 16 | 81 | **0.228** | **0.052** | **<0.001** |

Both in the total trend in abundance and in the proportion of species with declines, these results are similar to three European butterfly monitoring schemes (Table 3). Although the longer-running programs show larger cumulative declines, the annual rate of change in total abundance ranges from -2.0% to -2.6% for Ohio, Catalonia, and the Netherlands. The United Kingdom total abundance trends are split between generalist species (-0.8%) and specialist species (-2.4%). Across monitoring programs, declining species outnumber increasing species by a factor of two to three (Table 3).

**Table 3:** Comparison of this study’s results to European monitoring programs for rates of change in total abundance and classification of species trends as positive or negative. Number of sites represents those reported to contribute to the analysis, but may no longer be active. Number of butterflies counted per year is an approximation based on the most recent years of monitoring described in the references.

| Region (km<sup>2</sup>) | Years | Sites | Counted/year<br>(x 1000) | Annualized trend in total<br>abundance (cumulative) | Species' trends |  |  | Reference |
| --- | --- | --- | --- | --- | --- | --- | --- | --- |
|  |  |  |  |  | Positive | Negative | Stable/<br>not signif. |  |
| United Kingdom<br>(242,500) | 41 (1976-2017) | 3,164 | 1,700 | -0.8% (-28%) countryside<br>-2.4% (-63%) specialist | 11 | 22 | 24 | (51) |
| Netherlands<br>(42,508) | 25 (1992-2017) | 600 | 250 | -2.0% (-40%) | 11 | 23 | 13 | (52) |
| Catalonia, Spain<br>(32,108) | 22 (1994-2016) | 116 | 122 | -2.6% (-44%) | 15 | 46 | 5 | (40,53) |
| **Ohio, USA**<br>**(116,100)** | **20 (1996-2016)** | **104** | **80** | **-2.0% (-33%)** | **9** | **32** | **40** | **this study** |

In general, traits associated with species’ responses to climate were more important, based on the predictive ability (adjusted *R*^2^) of univariate models, than traits associated with habitat and host plant restrictions (Fig 5, Tables A and B in S1 Appendix). Phylogenetic signal was included for most traits’ models, so we focus on the phylogenetic generalized least squares results. The Monarch (*Danaus plexippus*) was the only migratory species in decline, although the others had erratic annual fluctuations that make trend estimation difficult (Fig 3). Species with more northern geographic ranges were associated with more negative population trends.

**Fig 5:**
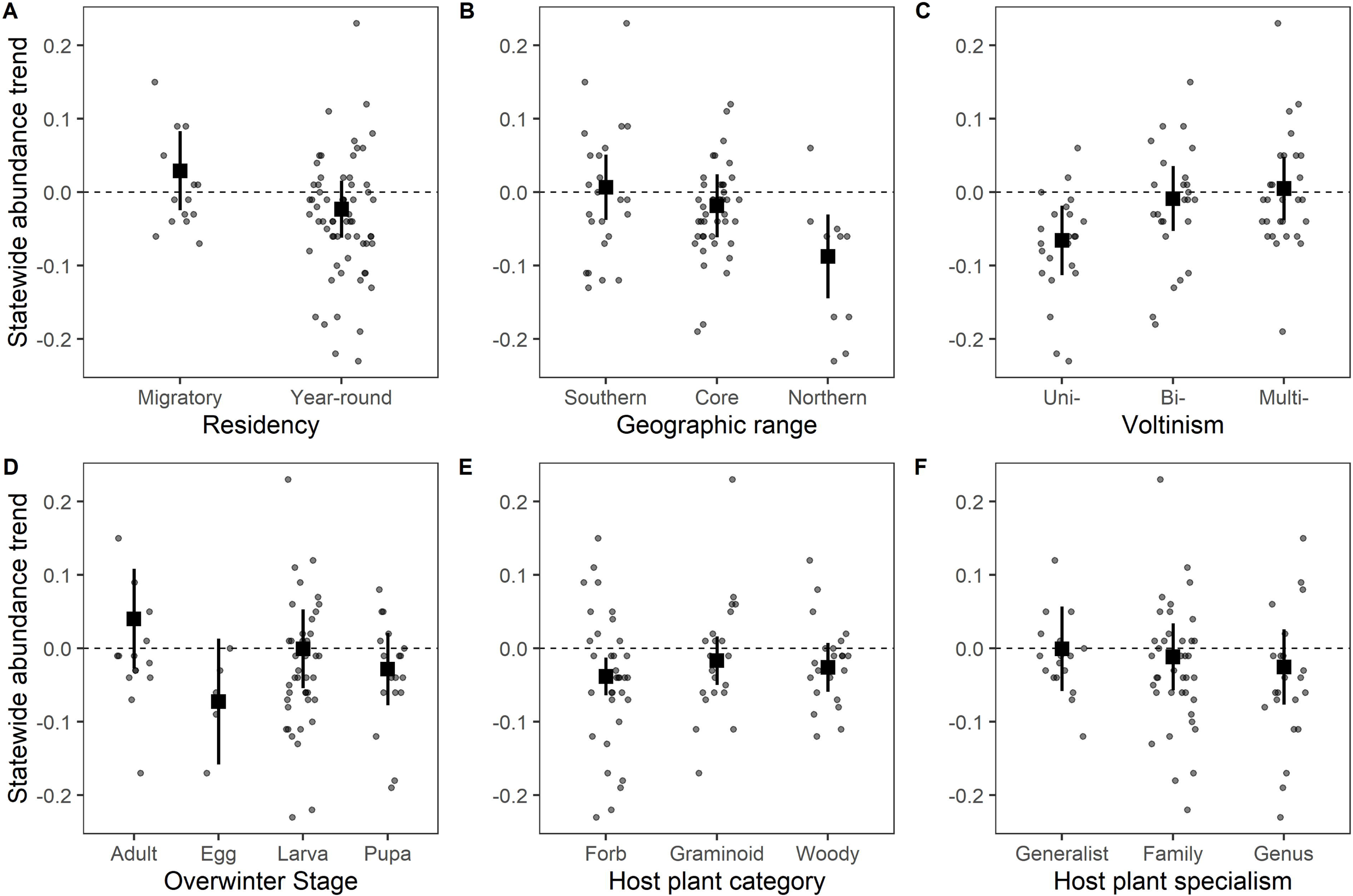
Species’ traits are associated with variation in the statewide trends in abundance. We plot each species’ trend compared to the six most important traits for the 78 species included in the phylogenetic GLS models with full results in Table A in S1 Appendix. Squares represent the regression coefficients with 95% confidence intervals shown in lines. Dots show trend estimates for each species from Table 1 uncorrected for phylogeny, jittered for visualization.

Univoltine species had more negative population trends than bivoltine or multivoltine species. Overwintering stage did not have a strong effect on trend. Species eating forb host plants had negative trends on average, but there was no effect of host plant specialization on population trends. Wing length, wetland habitat preference, or human-disturbed habitat preference were not associated with trends.

Our choice of modeling approach did not change the overall evidence of defaunation. Generalized linear mixed models with Poisson-distributed errors and generalized additive mixed models estimated declines in total abundance similar in magnitude at -1.83% and -2.13% annual rates, respectively. The annual trend estimate from TRIM, without sampling covariates, was half the magnitude at -0.96%. Species’ trends had high correlations between pairwise comparisons, but TRIM models estimated notably more positive trends compared to the other three approaches (Table C in S1 Appendix).

## Discussion

We show that the total butterfly abundance has declined by 33% over 20 years in Ohio. This rate is faster than the global abundance trend estimated for Lepidoptera (35% over 40 years) and corresponds more closely to the steeper declines (45% over 40 years) estimated for all insects (1). The Ohio butterfly monitoring program, judged by the weekly frequency, 20-year time period, and statewide spatial extent of its surveys, is the most extensive systematic insect survey in North America and comparable to three exemplary European butterfly monitoring schemes. The annualized 2% rate of decline in this study aligns closely with trends from European butterfly monitoring, confirming the decline of the most closely monitored group of insects in both Europe and North America (Table 3). With less known about other insect taxa, butterflies provide a necessary, if imperfect, surrogate to understand the trajectory and potential mechanisms behind broader insect trends (13). Extensive in both time and space, the decline in butterfly abundance reported here is the best estimate for the current rate of insect defaunation in North America.

The proportion of butterfly species with population declines compared to population increases is similar between Ohio (negative trends three times more numerous) and European studies (negative trends 2-3 times more numerous) (Table 3). In other taxa, moths in the United Kingdom show a similar proportion of species declines (68). Long-term monitoring in protected areas, although less extensive in space, shows more positive species trends for moths in Finland (at 67.7° latitude) and across pollinators in Spain (at 850-1750 m. elevations) (69,70). These counterexamples show how insect communities may shift at high-latitude or high-elevation sites with anthropogenic climate warming (21) or may persist in more remote areas. However, butterfly monitoring in populated areas show a consistency in observed declines (Table 3) that we argue would generalize to other landscapes dominated by human use.

We demonstrate abundance declines in species that are generalist, widespread, and not considered vulnerable to extinction (25,71). Although few may share concern for the most widespread, invasive butterfly in the world’s agricultural and urban settings (72), declines in *Pieris rapae* could be indicative of persistent environmental stressors that would affect other species as well. Generalist species that exploit human-disturbed habitat with annual rates of decline of more than 5% include *Lycaena phlaeas*, *Thymelicus lineola* (non-native), *Cercyonis pegala*, and *Colias eurytheme* (Table 2, Fig 4). We would expect negative environmental changes to disproportionately affect rare species prone to the demographic dangers of small populations or specialist species that rely on a narrow range of resources or habitat (UKBMS in Table 3, (24)). This pattern of species declines would lead to biotic homogenization as rarer species are lost and common, disturbance-tolerant species remain (73,74). However, our study adds another example of declines in common butterfly species thought to be well-suited to human-modified habitat (11,21,75).

The Eastern North American migratory Monarch (*Danaus plexippus*) abundance in Ohio is declining by 7% per year. The Monarch is the only declining migratory species out of 14 in our analysis. Despite disagreements about whether summer abundance trends have tracked winter colony declines (76,77), our study shows that the long-term trends correspond. However, our study’s first two years have very high Monarch population indices which could be outliers (Fig 3) following the two largest recorded winter population counts (16,78). With these two years removed, the statewide Monarch trend is a 4% decline per year, showing that the magnitude of summer abundance trends are sensitive to the years of data included. Our results align with a study using Illinois systematic monitoring data that shows a summer abundance decline for monarchs over two decades, but only during the period from 1994-2003, not from 2004-2013(79). A more recent study showed no decline during the summer during 2004-2016 using a population index from NABA counts (78). The trend we document comes from the sum of multiple summer breeding generations and fall migratory butterflies returning to Mexico; estimates of abundance for these separate generations may be required to model how different stages of the lifecycle contribute to the long-term decline in the winter colonies (78).

Our statewide analysis has potential limitations when used to evaluate individual species for potential conservation interventions or forecasts of population trajectories. Even with systematic monitoring, accurate estimates of insect abundance are missing from many species—a fifth of regularly observed species in Ohio did not meet our minimum data requirements to for us to estimate trends. None of these species are considered to be of conservation concern, but this also means that we would be limited in our ability to determine if their populations have reached threatened status. Targeted surveys of selected species, non-adult life stages, or rarely-sampled habitats can expand the monitoring to data-deficient species commonly excluded by protocols designed to monitor many species efficiently (51) and can be used to estimate demographic responses to environmental drivers not apparent from adult butterfly counts (80). Additional targeted species assessments could inform how worried we should be about the extreme population declines estimated for species observed at fewer than 10 monitoring sites (Table 2).

However, more data and more complex population models may not always lead to accurate predictions for insect population trajectories (81). Rather than recommending other systematic monitoring programs accumulate decades of data before assessing insect declines, we would advocate sharing data across regional programs to increase statistical power, as in (11), and integrating systematic monitoring with historical records and opportunistic observations to assess insect vulnerability more rapidly by using all potential sources of data (82,83).

Insect declines have multifaceted causes, and the relative impact of these causes is still unknown (84). Although analysis of the causes of site differences in abundance or species trends is beyond the scope of this study, we discuss three environmental drivers commonly associated with global insect declines: habitat loss and fragmentation, climate change, and agricultural intensification (84,85). If species’ traits are associated with population trends, then their relationships may suggest which environmental changes affect population responses in species sharing these traits (47,84,86). In this study, life-history traits were weakly predictive of population trends, but their associations provide hypotheses that could be tested further (47).

### Habitat loss and fragmentation

In Ohio, habitat loss and fragmentation plateaued well before butterfly monitoring started, with human population growth slowing by 1970. In common with other Midwestern states, Ohio had already lost tallgrass prairie species, such as the Regal Fritillary (*Speyeria idalia*), due to habitat conversion to agriculture (25,26). Land-use has changed slowly over the course of the monitoring program; fewer than 10% of monitoring sites have had more than 2.5% change in the surrounding (2-km radius) developed, agriculture, or forest land cover from 2001-2011 (29). The persistence of butterfly populations in a landscape of habitat fragments are mediated by species’ traits that permit them to either move between more isolated resources or persist in smaller, localized populations (85,87). Wing size is one life history trait associated with dispersal ability, but it had no association with species’ population trends (Tables A and B in S1 Appendix). However, defining habitat patches by land-use classes overlooks how mobile insect populations are bound by resources, varying across the lifecycle, rather than area (88,89). Although there has been little wholesale habitat conversion around our study transects, degradation of the remaining habitat could be a cause of the general decline in butterfly abundance.

### Climate change

Species trends are associated with two life-history traits, voltinism and range distribution, which suggest that the butterfly community is changing with the warming climate. Species that only complete one annual generation, or univoltine species, had more negative abundance trends. This aligns with obligate univoltine species becoming less common in Massachusetts (20), but is the opposite of the findings in Spain where multivoltine species are in steeper declines with exposure to increasingly dry summers (40). Multivoltine species may be more adaptive to annual and spatial variation in growing season length as many have plasticity in the voltinism observed within Ohio (25). For many species with flexible voltinism in Ohio, adding an extra generation in warmer summers increases their annual population growth rates (55). Northern-distributed species have more negative population trends compared to widely distributed or southern species. This corresponds with findings from Massachusetts and Europe that warm-adapted species are replacing cool-adapted species as range distributions shift (20,90). Even though these two traits should increase abundance for some species as the climate warms, it has not been enough to prevent the overall decline in butterfly abundance.

### Agricultural intensification

Cropland and pasture make up half of Ohio’s land area, so we would expect agricultural practices to affect statewide insect abundance. One assessment of pollinator habitat suitability based on land-use, conservation reserve program acreage, and crop type estimated an increase in resources in Ohio from 1982 through 2002, followed by a stable trend (91). However, agricultural practices can decrease insect abundance with systemic insecticides, herbicide use on host plants or nectar resources, and nitrogen fertilization that alters the composition of surrounding plant communities.

In Ohio, the use of neonicotinoids rapidly increased after 2004 when they became widely used on corn and soybeans (92,93). The mechanistic link between neonicotinoid insecticides and insect declines is established and observational studies have shown widespread impacts of their use (94–96). Even though seed-coatings with neonicotinoids reduce broadcast spraying, the mechanical planting of these seeds exposes widespread areas around farms to contaminated dust that exposes non-target plants and insects to biologically-relevant concentrations (97,98). In the United Kingdom and California, neonicotinoids are associated with butterfly declines (22,99) and hinder butterfly larval development on host plants (100). We did not design this study to test whether neonicotinoids affect butterfly abundance in Ohio. However, the observed declines across common and generalist species, which we otherwise would expect to exploit an agricultural or human-altered landscape, would be consistent with widespread exposure to insecticides.

Species that eat forbs as larvae have negative population trends (Fig 5). Both herbicide use and nitrogen deposition may alter plant communities to favor grasses over forbs (101). In Ohio, glyphosate use has increased linearly, and is now applied at 6 times the rate it was in 1996 (92,93). Milkweed losses, attributed to increased glyphosate use in the Midwest, contribute to declines in Monarch butterfly abundance (79,80). Nitrogen increases, which may come from fertilization or atmospheric deposition, have been linked to declines in grassland butterfly species adapted to low-nitrogen environments (102–104) and to higher mortality during larval development on enriched host plants (105).

## Conclusions

Systematic, long-term surveys of butterflies provide the most rigorous estimate for the rate of insect declines. This study demonstrates that defaunation is happening in North America similarly to Europe. In landscapes comprising natural areas amid heavy human land-use, butterfly total abundance is declining at 2% per year and 2-3 times more species have population trends declining rather than increasing. Additional Pollard-based monitoring programs in North America, listed in (9), will enable tracking insect trends over larger spatial extents as will efforts to integrate data across European monitoring schemes (11). The rates for other insect groups may deviate from this baseline and were previously estimated to be declining more rapidly than Lepidoptera (1). Expanded monitoring and support for taxonomists are imperative for other taxa and under sampled regions, like the Tropics where most insect diversity resides. Besides the evaluation if butterfly trends generalize to other insects, the most urgent research needs are understanding the causes of decline and testing mitigation strategies. As butterflies are the best-monitored insect taxa, they are the best indicator of the baseline threat to the 5.5 million insect species, the most diverse group of animals on earth.

## Acknowledgments

We thank the volunteers and directors who contribute their time and expertise to the Ohio butterfly monitoring program. The Ohio Department of Natural Resources provides support, the Cleveland Museum of Natural History provides data entry and archiving, and the Ohio Lepidopterists provides training and coordination for the Ohio butterfly monitoring program. We thank Marjorie Weber for advice on our phylogenetic analysis. The Department of the Interior Southeast Climate Adaptation Science Center and North Carolina State University supported TW during earlier work with Ohio butterflies that grew into this analysis.

---

**S1 Appendix. Supplementary methods and results.** Includes detailed methods for phenology models and phylogenetic trees, a figure of species trends plotted on a cladogram, two tables of model results from the trait analysis, and a table comparing our trend estimates with three other approaches.

## Notes

#### Summary of Updates

Revisions based on reviewer comments. New figures of all species population trends.

