## Supplementary material for "Butterfly abundance declines over 20 years of systematic monitoring in Ohio, USA": S1 Appendix

Phenology models

We used accumulated degree-days to account for the fact that development rate and phenological patterns of adult flight periods are temperature dependent. Daily maximum and minimum temperature data interpolated in gridded Daymet products were extracted for each site (28). These were used to estimate daily degree-days for each site using physiological thresholds of 5/30 C assuming a linear relationship between development rate and temperature within these bounds. Generic thresholds were used for all species as most do not have laboratory-derived physiological data (103). The lower threshold of 5C was chosen over a commonly-used threshold of 10C because the beginning of the monitoring season often started before any degree-days (10C base) had accumulated. We estimated daily degree-days for each site with an hourly interpolation method, based on the Daymet daily maximum and minimum temperatures, calibrated with hourly temperature data from OARDC weather stations in Ohio to account for daily temperature fluctuations due to sunrise and sunset timing across the year (35). Sites were clustered into four similarly-sized regions for phenology models and visualization (NE, NW, CN, and SW) based on geographic coordinates (104).

Similar to the UKBMS methods, population indices must have a weekly estimate of counts for each monitored site. Missing weeks are imputed using a generalized additive model that accounts for each species’ phenological patterns and variation in expected counts due to site differences in annual phenology and population size (31). We added geographic coordinates to account for spatial correlation between sites and climate gradients across the state and interactions to allow for regional variation in the season length constrained by calendar date (34). Random effects for site, year, and site x year accounted for variation in relative abundance between these factors. The error distribution was modeled as both Poisson and negative-binomial and the best model was selected by AIC for each species. The negative binomial error distribution allows for over-dispersion in the counts, which is common in insects due to fluctuations in phenology, clustering of individuals around floral resources, and high sensitivity to weather conditions. We fit models using the ‘bam’ function in the *mgcv* R package (105).

Phylogenetic analysis

We obtained coding sequences (CDS) for the most widely used DNA barcoding locus, the mitochondrial cytochrome c oxidase I gene COI-5P, from GenBank (www.ncbi.nlm.nih.gov/genbank/). For all species not found in GenBank, we obtained CDS from The Barcode of Life Data System (http://boldsystems.org/). When possible, we obtained sequences from multiple sampling locations in North America. Using custom python scripts, we translated these nucleotide sequences to sequences of peptides, and then we aligned all sequences in peptide space using MUSCLE v3.8.31 with default parameters (106). We removed regions of poor alignment with Gblocks v0.91b (107) using the options: -t=c -b1= “$b1” -b2= “$b1” -b3 = 1 -b4 = 6 -b5 = h, where b1 represents 70% of sampled sequences. For species with more than one sampling location, we took the consensus sequence from the filtered sequence alignment of all sampling locations using EMBOSS cons v.6.5.7 [default parameters]. This reduced our final alignment to 105 chromosomes, representing 104 species of butterflies in addition to the outgroup, the silkworm *Bombyx mori*. We visually inspected all alignments prior to phylogenetic reconstruction.

Owing to the relatively small size of our multiple-species alignment—i.e. a single mtDNA locus, 651 bp in length—we decided to take both a constrained and unconstrained maximum likelihood approach to estimate the genealogical relationships of our samples. Our unconstrained approach represents the raw gene genealogy for COI-5P across all 105 species. We used RAxML v8.2.12 [options: -f a -m GTRGAMMA -x 12345 -p 12345 -# 10000] to generate this gene tree and perform bootstrapping (108). Some of the species from our analysis, though not all, were recently used in a more comprehensive phylogenetic analysis of butterflies (59), thus prompting us to constrain the phylogenetic backbone of our tree using family-level relationships. Using a custom python script, we generated a multi-furcating constraint tree, whereby species were grouped together at the family level. We again used RAxML to generate a constrained tree using the options: -f a -m GTRGAMMA -x 12345 -p 12345 -# 10000 -g backboneConstraint.txt. Both trees were re-rooted by *Bombyx mori*, and all re-rooting and cladograms was generated using FigTree v1.4.4 (https://github.com/rambaut/figtree). All accessions, sequence alignments, and phylogenetic relationships are deposited in Dryad.

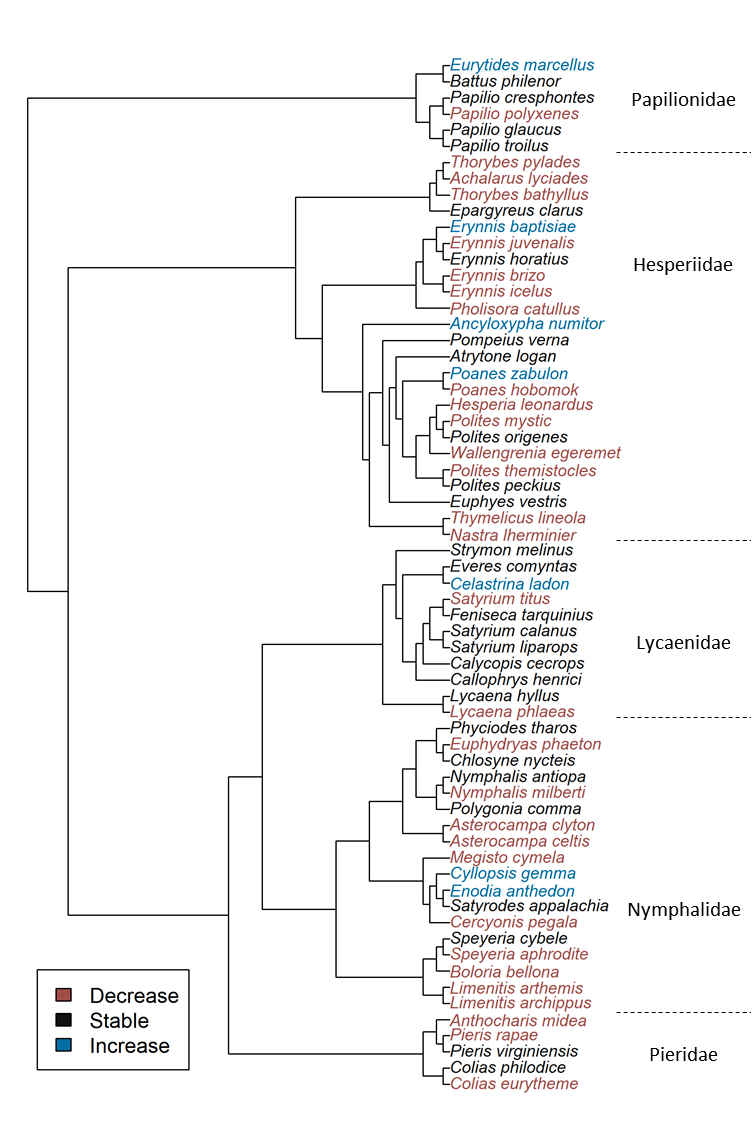

Fig A: Species’ abundance trend categories plotted over a cladogram constrained by the labeled families with estimated branch lengths ignored for visualization. Some genera were not monophyletic as our analysis was limited to one mtDNA locus.

Table A: Analysis of life-history traits associated with species abundance trends across 81 species. Each trait was modeled as a univariate predictor of species trends.

|  | Linear models | | | | |  | Phylogenetic generalized least squares | | | | | |
| --- | --- | --- | --- | --- | --- | --- | --- | --- | --- | --- | --- | --- |
| Covariate | Coef. | Std.error | *t* | *P* | Adj. R^2^ |  | Coef. | Std.error | *t* | *P* | Adj. R^2^ | Pagel's λ |
| Migratory | 0.01 | 0.02 | 0.38 | 0.71 | 0.12 |  | 0.03 | 0.03 | 1.06 | 0.29 | 0.05 | 0.47 |
| Resident | -0.03 | 0.01 | -3.65 | <0.01 |  |  | -0.02 | 0.02 | -1.17 | 0.25 |  |  |
| Range: northern | -0.08 | 0.02 | -3.78 | <0.001 | 0.17 |  | -0.09 | 0.03 | -3.02 | <0.01 | 0.13 | 0.56 |
| Range: core | -0.03 | 0.01 | -2.15 | 0.03 |  |  | -0.02 | 0.02 | -0.86 | 0.39 |  |  |
| Range: southern | -0.01 | 0.02 | -0.39 | 0.70 |  |  | 0.01 | 0.02 | 0.29 | 0.78 |  |  |
| Univoltine | -0.07 | 0.02 | -4.34 | <0.01 | 0.18 |  | -0.07 | 0.02 | -2.73 | <0.01 | 0.14 | 0.54 |
| Bivoltine | -0.02 | 0.02 | -1.14 | 0.26 |  |  | -0.01 | 0.02 | -0.39 | 0.70 |  |  |
| Multivoltine | -0.01 | 0.02 | -0.34 | 0.74 |  |  | 0.00 | 0.02 | 0.22 | 0.83 |  |  |
| Host type: forb | -0.04 | 0.01 | -3.00 | <0.01 | 0.10 |  | -0.04 | 0.01 | -2.91 | <0.01 | -0.01 | 0.00 |
| Host type: graminoid | -0.01 | 0.02 | -0.64 | 0.52 |  |  | -0.02 | 0.02 | -1.01 | 0.32 |  |  |
| Host type: woody | -0.02 | 0.02 | -1.49 | 0.14 |  |  | -0.03 | 0.02 | -1.54 | 0.13 |  |  |
| Host genus-specific | -0.04 | 0.02 | -2.67 | <0.01 | 0.09 |  | -0.03 | 0.03 | -0.97 | 0.34 | -0.01 | 0.55 |
| Host family-specific | -0.03 | 0.01 | -1.99 | 0.05 |  |  | -0.01 | 0.02 | -0.50 | 0.62 |  |  |
| Host generalist | -0.01 | 0.02 | -0.58 | 0.57 |  |  | 0.00 | 0.03 | -0.02 | 0.99 |  |  |
| Winter stage: adult | -0.01 | 0.02 | -0.44 | 0.67 | 0.09 |  | 0.04 | 0.04 | 1.16 | 0.25 | 0.05 | 0.62 |
| Winter stage: egg | -0.06 | 0.03 | -1.89 | 0.06 |  |  | -0.07 | 0.04 | -1.66 | 0.10 |  |  |
| Winter stage: larva | -0.02 | 0.01 | -1.95 | 0.06 |  |  | 0.00 | 0.03 | -0.03 | 0.97 |  |  |
| Winter stage: pupa | -0.04 | 0.02 | -2.07 | 0.04 |  |  | -0.03 | 0.03 | -1.14 | 0.26 |  |  |
| Wing size | 0.00 | 0.01 | -0.06 | 0.96 | -0.01 |  | 0.00 | 0.01 | 0.30 | 0.77 | -0.01 | 0.50 |
| Wetland habitat specialists | 0.01 | 0.03 | 0.36 | 0.72 | -0.01 |  | 0.00 | 0.03 | -0.07 | 0.94 | -0.01 | 0.50 |
| Disturbed habitat generalists | 0.02 | 0.02 | 1.29 | 0.20 | 0.01 |  | 0.03 | 0.02 | 1.69 | 0.10 | 0.02 | 0.58 |

Table B: Analysis of life-history traits associated with species abundance trends across 67 non-migratory species. Each trait was modeled as a univariate predictor of species trends.

|  | Linear models | | | | |  | Phylogenetic generalized least squares | | | | | |
| --- | --- | --- | --- | --- | --- | --- | --- | --- | --- | --- | --- | --- |
| Covariate | Coef. | Std.error | *t* | *P* | Adj. R^2^ |  | Coef. | Std.error | *t* | *P* | Adj. R^2^ | Pagel's λ |
| Range: northern | -0.08 | 0.02 | -3.67 | <0.001 | 0.19 |  | -0.09 | 0.03 | -3.11 | 0.00 | 0.10 | 0.50 |
| Range: core | -0.02 | 0.01 | -1.92 | 0.06 |  |  | -0.02 | 0.02 | -0.97 | 0.34 |  |  |
| Range: southern | -0.02 | 0.02 | -1.26 | 0.21 |  |  | -0.01 | 0.02 | -0.54 | 0.59 |  |  |
| Univoltine | -0.07 | 0.02 | -4.29 | <0.01 | 0.23 |  | -0.07 | 0.02 | -3.01 | <0.01 | 0.15 | 0.52 |
| Bivoltine | -0.04 | -0.02 | -2.05 | 0.04 |  |  | -0.04 | 0.02 | -1.50 | 0.14 |  |  |
| Multivoltine | -0.00 | 0.02 | -0.23 | 0.82 |  |  | 0.00 | 0.02 | 0.17 | 0.86 |  |  |
| Host type: forb | -0.06 | 0.02 | -4.11 | <0.001 | 0.20 |  | -0.06 | 0.02 | -4.14 | <0.001 | 0.04 | 0.00 |
| Host type: graminoid | -0.01 | 0.02 | -0.54 | 0.59 |  |  | -0.02 | 0.02 | -0.90 | 0.37 |  |  |
| Host type: woody | -0.03 | 0.02 | -1.47 | 0.15 |  |  | -0.03 | 0.02 | -1.54 | 0.13 |  |  |
| Host genus-specific | -0.06 | 0.02 | -3.36 | <0.01 | 0.16 |  | -0.05 | 0.03 | -1.96 | 0.06 | 0.00 | 0.42 |
| Host family-specific | -0.03 | 0.01 | -2.14 | 0.04 |  |  | -0.02 | 0.02 | -1.06 | 0.30 |  |  |
| Host generalist | -0.01 | 0.02 | -0.56 | 0.58 |  |  | -0.01 | 0.03 | -0.46 | 0.65 |  |  |
| Winter stage: adult | -0.07 | 0.05 | -1.41 | 0.16 | 0.13 |  | -0.04 | 0.06 | -0.65 | 0.52 | -0.01 | 0.45 |
| Winter stage: egg | -0.06 | 0.03 | -1.85 | 0.07 |  |  | -0.08 | 0.04 | -1.89 | 0.06 |  |  |
| Winter stage: larva | -0.03 | 0.01 | -2.06 | 0.04 |  |  | -0.02 | 0.03 | -0.66 | 0.51 |  |  |
| Winter stage: pupa | -0.04 | 0.02 | -2.02 | 0.05 |  |  | -0.03 | 0.02 | -1.27 | 0.21 |  |  |
| Wing size | 0.00 | 0.01 | -0.01 | 0.99 | -0.02 |  | 0.00 | 0.01 | 0.41 | 0.68 | -0.01 | 0.39 |
| Wetland habitat specialists | 0.02 | 0.03 | 0.67 | 0.50 | -0.01 |  | 0.01 | 0.03 | 0.27 | 0.79 | -0.02 | 0.38 |
| Disturbed habitat generalists | 0.01 | 0.02 | 0.39 | 0.70 | -0.01 |  | 0.01 | 0.02 | 0.69 | 0.49 | -0.01 | 0.47 |

Table C: Comparison of species trends classified as negative, stable, or positive between the modeling approach presented in the main text (generalized linear mixed model (GLMM) with negative binomial error distributions) versus three alternative models: GLMM with Poisson error distribution, generalized additive mixed model (GAMM) with negative binomial error distribution, and the generalized linear model with serial correlation and over-dispersion fit by TRIM. Pairwise Pearson correlations between 81 species trends between two modeling approaches are reported in the right column.

|  |  | **GLMM (negative binomial)** | | |  |  |
| --- | --- | --- | --- | --- | --- | --- |
|  |  | Negative | Stable | Positive | Total | R = 0.86 |
| **GLMM (poisson)** | Negative | 26 | 2 | 0 | 28 |  |
|  | Stable | 5 | 35 | 1 | 41 |  |
|  | Positive | 1 | 3 | 8 | 12 |  |
|  | Total | 32 | 40 | 9 | 81 |  |
|  |  | Negative | Stable | Positive | Total | R = 0.94 |
| **GAMM (negative binomial)** | Negative | 29 | 1 | 0 | 30 |  |
|  | Stable | 3 | 38 | 3 | 44 |  |
|  | Positive | 0 | 1 | 6 | 7 |  |
|  | Total | 32 | 40 | 9 | 81 |  |
|  |  | Negative | Stable | Positive | Total | R = 0.74 |
| **TRIM (over-dispersed poisson)** | Negative | 19 | 5 | 0 | 24 |  |
|  | Stable | 12 | 25 | 1 | 38 |  |
|  | Positive | 1 | 10 | 8 | 19 |  |
|  | Total | 32 | 40 | 9 | 81 |  |
